# Topical GZ21T inhibits the growth of actinic keratoses in a UVB induced model of skin carcinogenesis

**DOI:** 10.1101/2022.09.07.506864

**Authors:** Zachary A. Bordeaux, Justin Choi, Gabriella Braun, Cole Davis, Melika Marani, Kevin Lee, Christeen Samuel, Jackson Adams, Reed Windom, Anthony Pollizzi, Anusha Kambala, Hannah Cornman, Sriya V. Reddy, Weiying Lu, Olusola O. Oladipo, Martin P. Alphonse, Cameron E. West, Shawn G. Kwatra, Madan M. Kwatra

**Affiliations:** Department of Dermatology, Johns Hopkins University School of Medicine; Department of Anesthesiology, Duke University School of Medicine; Genzada Pharmaceuticals; US Dermatology Partners; Department of Oncology, Johns Hopkins University School of Medicine; Department of Pharmacology and Cancer Biology, Duke University School of Medicine

**Keywords:** Actinic keratoses, Ultraviolet B, curcumin, harmine, isovanillin, carcinogenesis

## Abstract

Actinic keratoses (AKs) are premalignant intraepidermal neoplasms that occur as a result of cumulative sun damage. AKs commonly relapse, and up to 16% undergo malignant transformation into cutaneous squamous cell carcinoma (cSCC). There is a need for novel therapies that reduce the quantity and surface area of AKs as well as prevent malignant transformation to cSCCs. We recently showed that GZ17-6.02, an anti-cancer agent composed of curcumin, haramine, and isovanillin, inhibited the growth of H297.T cells. The present study evaluated the efficacy of a novel topical formulation of GZ17-6.02, known as GZ21T, in a murine model of AK generated by exposing SKH1 mice to ultraviolet irradiation. Treatment of mice with topical GZ21T inhibited the growth of AKs by decreasing both lesion count (*p*=.028) and surface area occupied by tumor (*p*=.026). GZ21T also suppressed the progression of AKs to cSCC by decreasing the count (*p*=.047) and surface area (*p*=.049) of lesions more likely to represent cSCC. RNA sequencing and proteomic analyses revealed that GZ21T suppressed several pathways, including MAPK (*p*=.026), Pi3K-Akt (*p*=.028), HIF-1α (*p*=.030), Wnt (*p*=.031), insulin (*p*=.011), and ErbB (*p*=.006) signaling. GZ21T also upregulated the autophagy-promoting protein AMPK, while suppressing proteins such as PD-L1, glutaminase, pAkt1 S473, and eEF2K.

**GRAPHICAL ABSTRACT:** 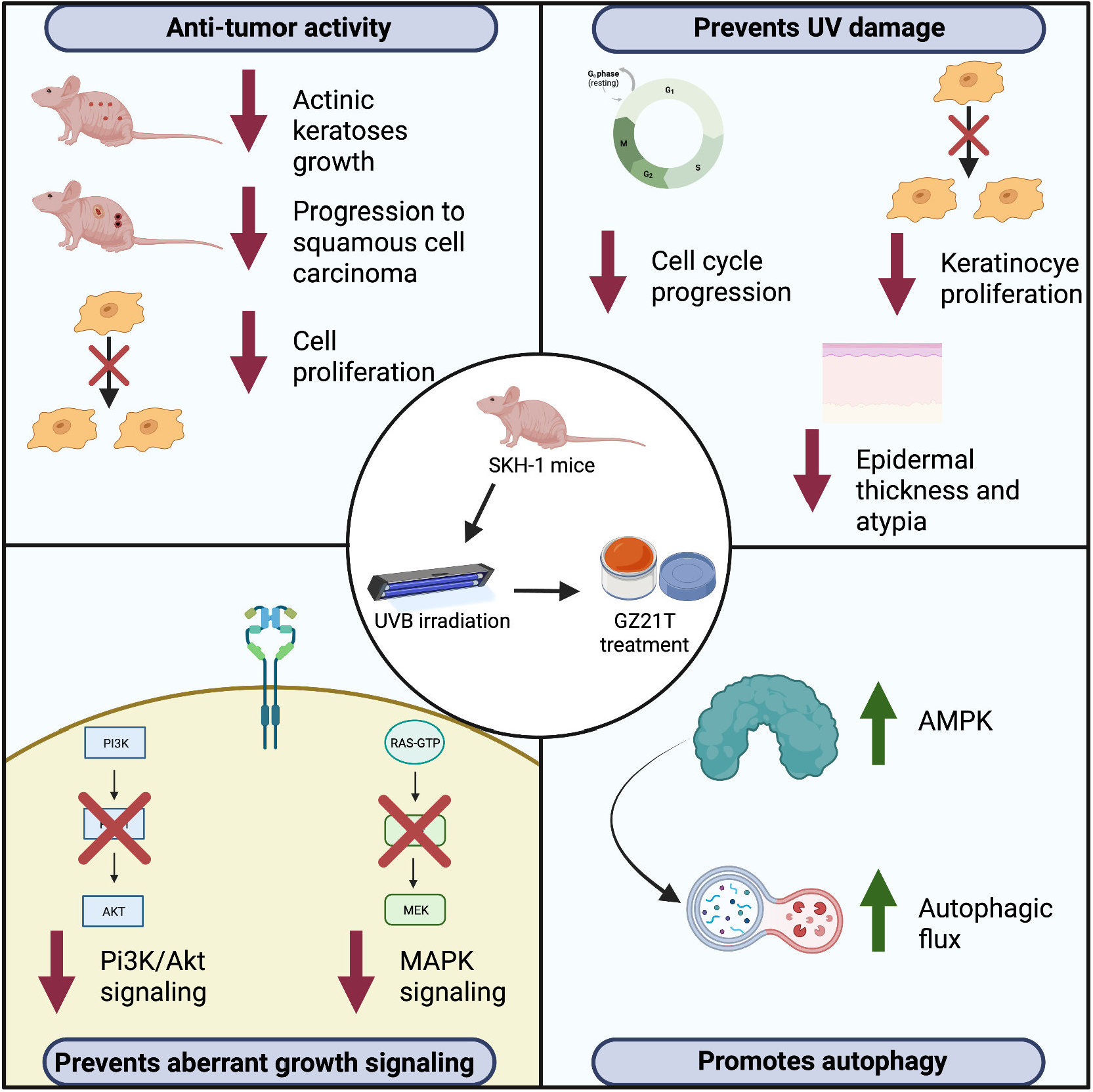

## INTRODUCTION

Actinic keratoses (AKs) are premalignant intraepidermal neoplasms characterized by keratinocyte atypia in the basal layer of the epidermis that carry an estimated prevalence of 28-49% (Berman & Cockerell, 2013; Fernandez Figueras, 2017; Flohil et al., 2013). AKs arise as a result of cumulative sun damage and present as rough, scaly patches, plaques, or papules on an erythematous base (Dirschka et al., 2017), typically affecting sun-exposed areas in older, fair-skinned individuals (Boone et al., 2016; Costa et al., 2015; Fernandez Figueras, 2017; Salmon & Tidman, 2016). Patients with few or isolated AKs are typically treated with lesion-directed therapies such as cryotherapy, laser therapy, or curettage, while patients with more diffuse lesions may require field-directed therapies such as 5-fluorouracil, diclofenac, imiquimod, or photodynamic therapy. Furthermore, it is estimated that up to 16% of AKs undergo malignant transformation into cutaneous squamous cell carcinoma (cSCC) (Dirschka et al., 2017). Given the side effect profiles and efficacy of current treatment options, novel therapeutic options for the treatment of AKs are needed (Arenberger & Arenbergerova, 2017; Taute et al., 2017).

The present study was undertaken to evaluate GZ12T, the topical formulation of GZ17-6.02, in a murine model of AK. The selection of GZ17-6.02, a novel anti-cancer agent composed of curcumin (10% by weight), isovanillin (77% by weight), and harmine (13% by weight) (Ghosh *et al.* 2019), was based on its activity against a variety of malignancies, including head and neck squamous cell carcinoma, pancreatic ductal adenocarcinoma, melanoma, and glioblastoma (Booth et al., 2022; Booth et al., 2021; Choi et al., 2022; Ghosh et al., 2019; Vishwakarma et al., 2018). Finally, our preliminary evaluation of GZ17-6.02’s activity against H279.T cells, a fibroblast cell line derived from an AK patient, demonstrated its efficacy via modulation of autophagy and cell death (West et al., 2022).

The antitumor activity of GZ17-6.02 appears to be due to the inherent synergism of its components. Curcumin is a well-studied anti-tumor agent found in the dietary spice turmeric that suppresses tumor progression through inhibition of pathways such as PI3K-AKT/mTOR signaling, which has previously been implicated in the pathogenesis of AK and cSCC (Maiti et al., 2019; Thomson et al., 2021; Vallianou et al., 2015). Curcumin has also been found to activate the tumor suppressor p53 and to suppress epidermal growth factor receptor (EGFR) signaling by blocking the binding of early growth response-1 (Egr-1) to the EGFR promoter and inhibiting the phosphorylation of EGFR (Chen et al., 2006; Sun et al., 2012; Vollono et al., 2019; Zhen et al., 2014). Similarly, harmine is shown to disrupt EGFR signaling by inhibiting the dual-specificity tyrosine-(Y)-phosphorylation-regulated kinase, DYRK1A (Pozo et al., 2013), and to inhibit Akt/mTOR and ERK signaling (Chamcheu et al., 2019). Harmine also exerts antitumor activity by inhibiting DNA replication, inducing DNA damage, reducing cell cycle progression, and suppressing angiogenesis (Zhang et al., 2020).

Here, we evaluate the efficacy of topical GZ12T against AK *in vivo* using an ultraviolet B (UVB) induced model of skin carcinogenesis. Our results show that GZ21T inhibits AK growth, blocks its progression to cSCC, and exerts a cytoprotective effect with short-term UVB exposure. The underlying mechanism, revealed by transcriptomic and proteomic analyses, involves its action at multiple levels, including the downregulation of pathways previously implicated in the pathogenesis of AK such as MAPK, Pi3K-Akt, Wnt, and ErbB signaling, as well as inhibition of proteins such as PD-L1, glutaminase, pAkt1 S473, and eEF2K (Brennan et al., 2013; Ratushny et al., 2012).

## RESULTS

### Topical GZ21T inhibits AK growth in a UVB-induced model of skin carcinogenesis

To assess the ability of topical GZ21T to inhibit AK growth *in vivo*, we used a UVB-induced model of skin carcinogenesis. An overview of the experimental design is shown in Figure 1A, while representative images and histology of AK lesions from control and GZ21T-treated mice after eight weeks of treatment are shown in Figure 1B and Figure 1C, respectively. After eight weeks of treatment, mice receiving GZ21T demonstrated a significant decrease in both lesion count (*p*=.028, Figure 1D) and surface area occupied by tumor (*p*=.026, Figure 1E).

**Figure 1:**
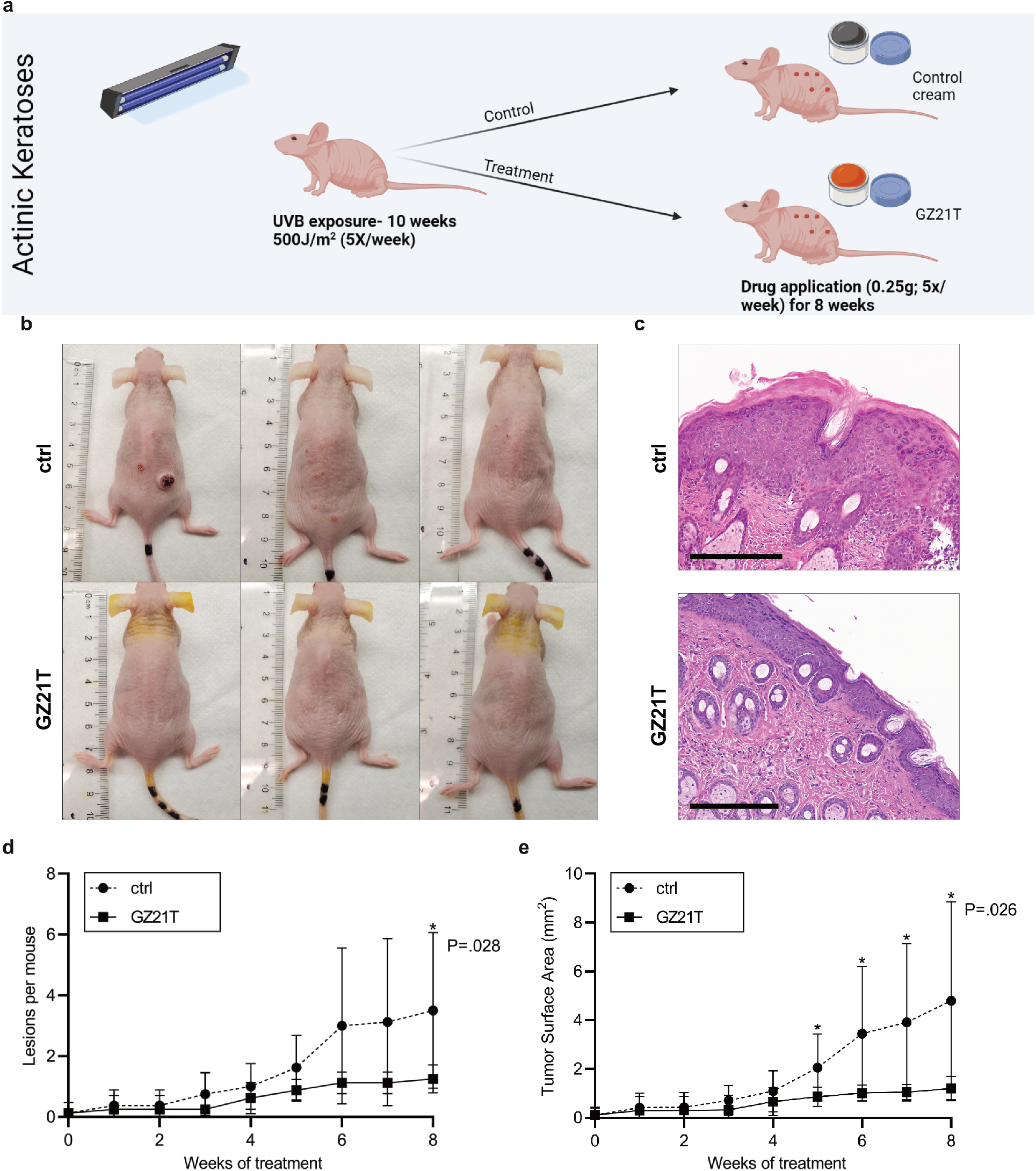
GZ21T inhibits the growth of actinic keratoses. A) To induce AK lesions, mice were irradiated with 500 J/m^2^ UVB five days per week for ten weeks and were subsequently treated with 0.25 g GZ21T or control cream five days per week for eight weeks. B) Representative images of control and GZ21T treated mice after eight weeks of treatment. C) Representative H&E staining of lesional samples from control and GZ21T treated mice after eight weeks of treatment. Black boxes represent 300 μm. D) Lesion count and E) tumor surface area over time for control and GZ21T treated mice. * P<0.05; ctrl, control.

### Molecular mechanism of AK progression

To better understand the targets that may be affected by GZ21T, we first performed transcriptomic and proteomic analyses to identify the molecular changes that occur with UVB exposure and progression from healthy skin to AK. Figure 2A provides a heatmap of the differentially expressed genes (DEGs) between lesional, peri-lesional (PL), non-lesional (NL), and non-UV exposed skin. Interestingly, progression from non-UV exposed skin to NL, PL, and lesional tissue resulted in an increasing number of DEGs. Compared to non-UV exposed skin, NL samples showed 194 DEGs (Figure 2B), PL tissue showed 731 DEGs (Figure 2C), and lesional skin 1502 DEGs (Figure 2D). Of the DEGs, 45 were unique to NL tissue, 208 were unique to PL skin, 994 were unique to lesional samples, and 114 were shared among lesional, PL, and NL skin (Figure 2E).

**Figure 2:**
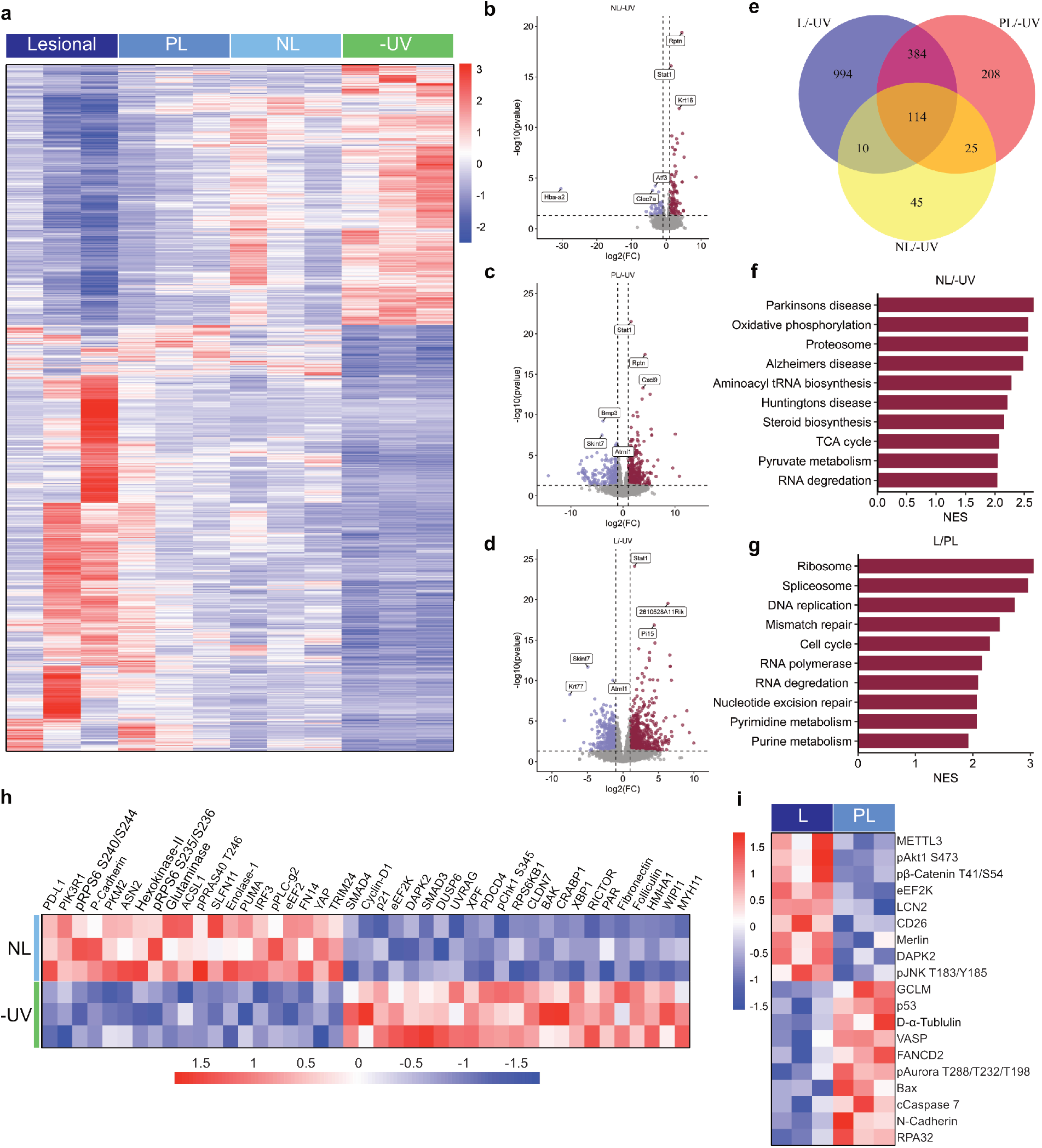
Molecular mechanism of actinic keratoses progression. A) Heatmap of DEGs in lesional, PL, NL, and non-UV exposed mouse skin. B) Volcano plots for DEGs between NL and non-UV exposed skin, C) PL and NL skin, and D) lesional and PL skin. E) Venn diagram showing unique and shared DEGs between lesional, PL, and NL skin compared to non-UV exposed skin. F) GSEA results for pathways enriched in NL compared to non-UV exposed skin and G) lesional compared to PL skin. H) Heatmap of differentially expressed proteins between NL and non-UV exposed skin. I) Differentially expressed proteins between lesional and PL skin. PL, peri-lesional; NL, non-lesional; -UV, non-UV exposed; FC, fold change; NES, normalized enrichment score; L, lesional.

We next conducted gene set enrichment analyses (GSEA) comparing NL to non-UV exposed skin and PL to lesional tissue to identify pathways altered by UV exposure and progression to AK. UV irradiation upregulated several metabolic and synthetic pathways such as oxidative phosphorylation, proteosome, steroid biosynthesis, and the TCA cycle (Figure 2F). Progression from PL skin to AK was accompanied by enrichment for pathways involved in DNA synthesis, protein synthesis, and cell cycle progression, such as ribosome, DNA replication, cell cycle, and purine and pyrimidine metabolism (Figure 2G).

Finally, we evaluated the proteomic alterations that occur with UV exposure and progression to AK using reverse phase protein arrays (RPPA). A comparison of NL and non-UV exposed skin showed an increase in 20 and a decrease in 23 proteins (Figure 2H). Notably, NL tissue was characterized by upregulation of the T-cell inhibitory protein PD-L1 and glutaminase. Comparison of lesional and PL tissue showed that progression to AK was accompanied by increased expression of 9 and decreased expression of 9 proteins (Figure 2I). Among the upregulated proteins were pAkt1 S473, a component of the Pi3K-Akt signaling pathway, and eEF2K, which supports the survival of neoplastic tissue under conditions of nutrient deprivation (Temme & Asquith, 2021).

### Transcriptomic effects of GZ21T treatment

To better understand the mechanism behind GZ21T’s anti-tumor activity, we next performed RNA sequencing on lesional skin from control and treated mice. Differential expression analysis revealed 133 DEGs, of which 99 were upregulated, and 54 were downregulated, in GZ21T treated mice (Figures 3A-B). GSEA of GZ21T-treated skin revealed downregulation of pathways involved in DNA synthesis, protein synthesis, metabolism, and cell cycle progression, such as DNA replication, ribosome, proteasome, oxidative phosphorylation, and cell cycle (Figure 3C). Interestingly, many of these pathways were upregulated by UV exposure and with progression from healthy skin to AK (Figure 2F), indicating that GZ21T may reverse the adverse effects of UV exposure. We next evaluated the effect of GZ21T on pathways previously implicated in the pathogenesis of AK using gene set variation analysis (GSVA) (Thomson et al., 2021). This analysis revealed suppression of MAPK (*p*=.026), Pi3K-Akt (*p*=.028), HIF-1α (*p*=.030), Wnt (*p*=.031), insulin (*p*=.011), and ErbB (*p*=.006) signaling pathways in lesional tissue of GZ21T-treated mice (Figure 3D). Of note, these pathways were not downregulated in NL skin, suggesting that GZ21T may preferentially suppress these targets in neoplastic tissue.

**Figure 3:**
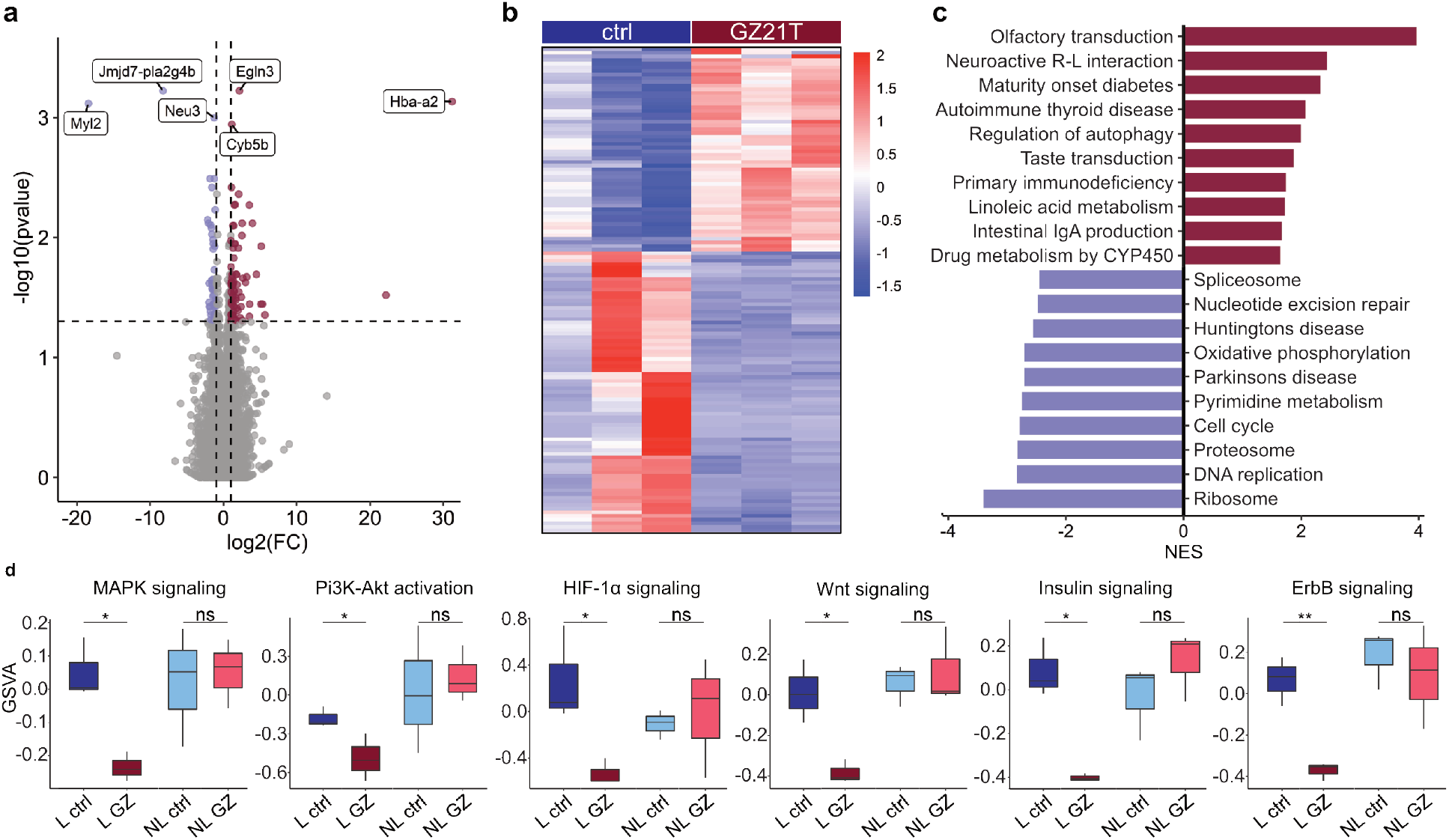
Transcriptomic analysis of control and GZ21T treated skin. A) Volcano plot and B) heatmap of DEGs between lesional samples of control and GZ21T treated mice. C) GSEA results showing pathways up and downregulated by GZ21T in lesional tissue. D) GSVA for pathways previously implicated in the pathogenesis of AK in lesional and non-lesional tissue from control and GZ21T treated mice. * P<0.05; ** P<0.01; FC, fold change; ctrl, control; R-L, receptor-ligand; NES, normalized enrichment score; GSVA, gene set variation analysis; L, lesional; NL, non-lesional; GZ, GZ21T; ns, not significant.

### Proteomic effects of GZ21T treatment

We also evaluated the proteins that were altered by GZ21T treatment using RPPA. Figure 4A shows a heatmap of the significantly different proteins between the lesional skin of control and treated mice, while Figure 4B shows the proteins with a change in expression greater than 25% following treatment. Interestingly, GZ21T modulated the expression of several proteins that were altered by UV exposure (Figure 2H). For example, PD-L1 and glutaminase were downregulated while PAR was upregulated. GZ21T also altered the expression of several proteins that we found to be involved in the progression from PL tissue to AK (Figure 2I). Levels of pAkt1 S473, eEF2K, and LCN2 were decreased while RPA32 and FANCD2 were increased. We also found that the autophagy promoting protein, AMPK, was upregulated by GZ21T (Li & Chen, 2019). Next, we performed EnrichR functional enrichment analysis on gene lists of proteins downregulated by GZ21T to confirm the alterations in pathways identified by our transcriptomic analysis. Similar to our RNA sequencing results, this analysis showed suppression of ErbB signaling, cell cycle, insulin signaling, HIF-1α signaling, Pi3K-Akt signaling, and MAPK signaling in GZ21T treated skin (Figure 4C). We also constructed a GeneMANIA network using gene names of proteins downregulated by GZ21T to highlight genes with shared protein domains and physical interactions or that are co-localized or co-expressed with the targets of GZ21T (Figure 4D). We found that the most over-represented functions of these proteins were related to peptidyl-serine modification, positive regulation of DNA metabolism, cell cycle checkpoints, DNA integrity checkpoints, G2/M phase transition, lymphocyte apoptosis, and the response to UV.

**Figure 4:**
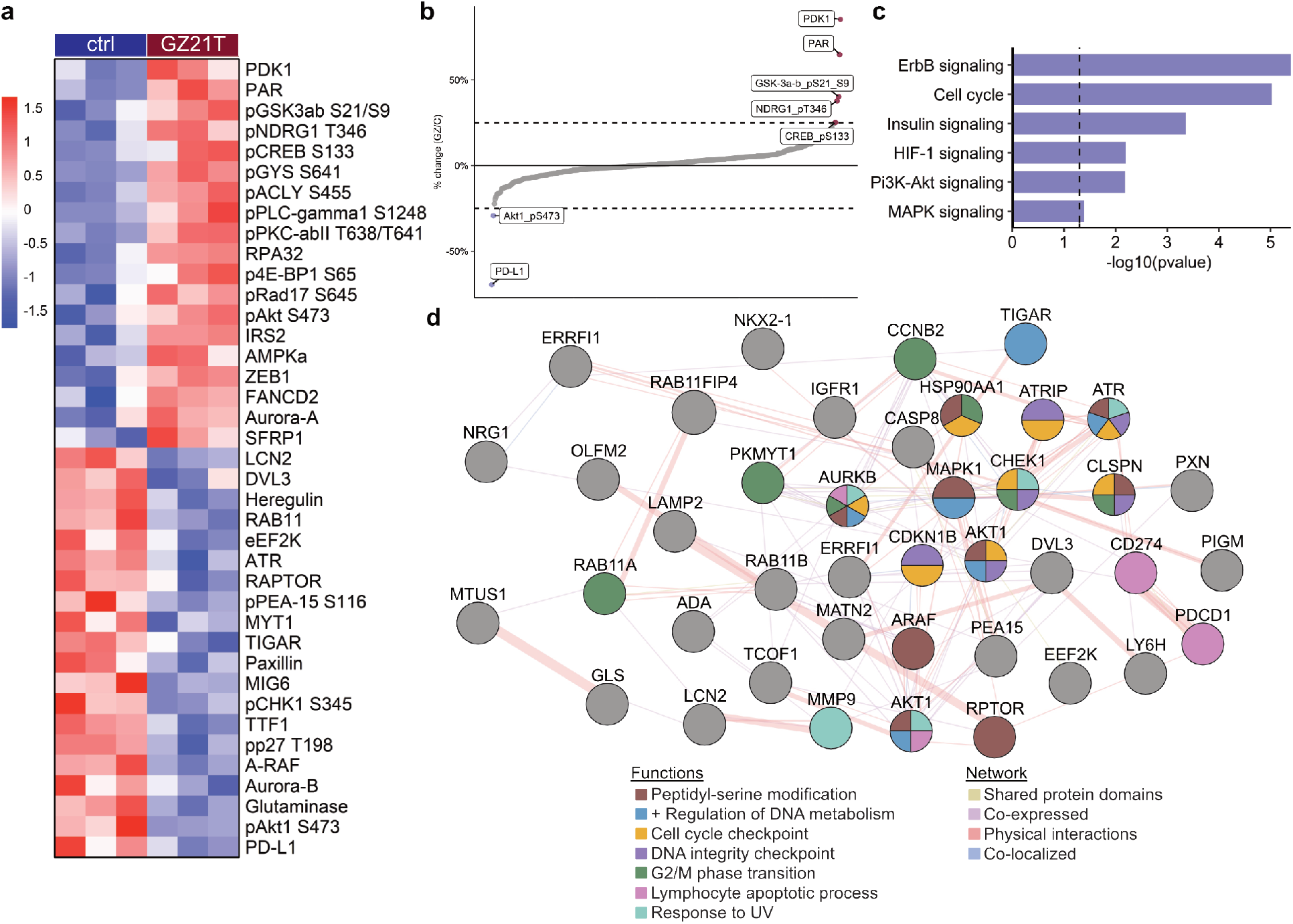
Proteomic analysis of control and GZ21T treated skin. A) Heatmap of differentially expressed proteins identified by RPPA between lesional samples of control and GZ21T treated mice. B) Significant proteins showing a percent change greater than 25% between lesional samples of control and GZ21T treated mice. C) EnrichR enrichment analysis of proteins downregulated by GZ21T in lesional samples for pathways implicated in the pathogenesis of AK. D) GeneMANIA network of gene lists of proteins downregulated by GZ21T in lesional AK skin. ctrl, control; UV, ultra violet.

### GZ21T prevents the progression of AK to cSCC

We next sought to determine if GZ21T could prevent the progression of AK to cSCC. To induce cSCC lesions, we prolonged the UV exposure period from ten weeks in the AK experiment to 14 weeks (Figure 5A). After six weeks of treatment, mice were analyzed for the presence of lesions greater than 3 mm in diameter, which are more likely to represent cSCC than AK (de Gruijl & van der Leun, 1991). Representative images of control and GZ21T treated mice after six weeks of treatment are shown in Figure 5B. Analyses of lesions greater than 3 mm in diameter demonstrated a significant decrease in lesion count (*p*=.047, Figure 5C) and surface area covered by tumor (*p*=.049, Figure 5D) in GZ21T treated mice, suggesting that GZ21T may inhibit the progression of AK to cSCC.

**Figure 5:**
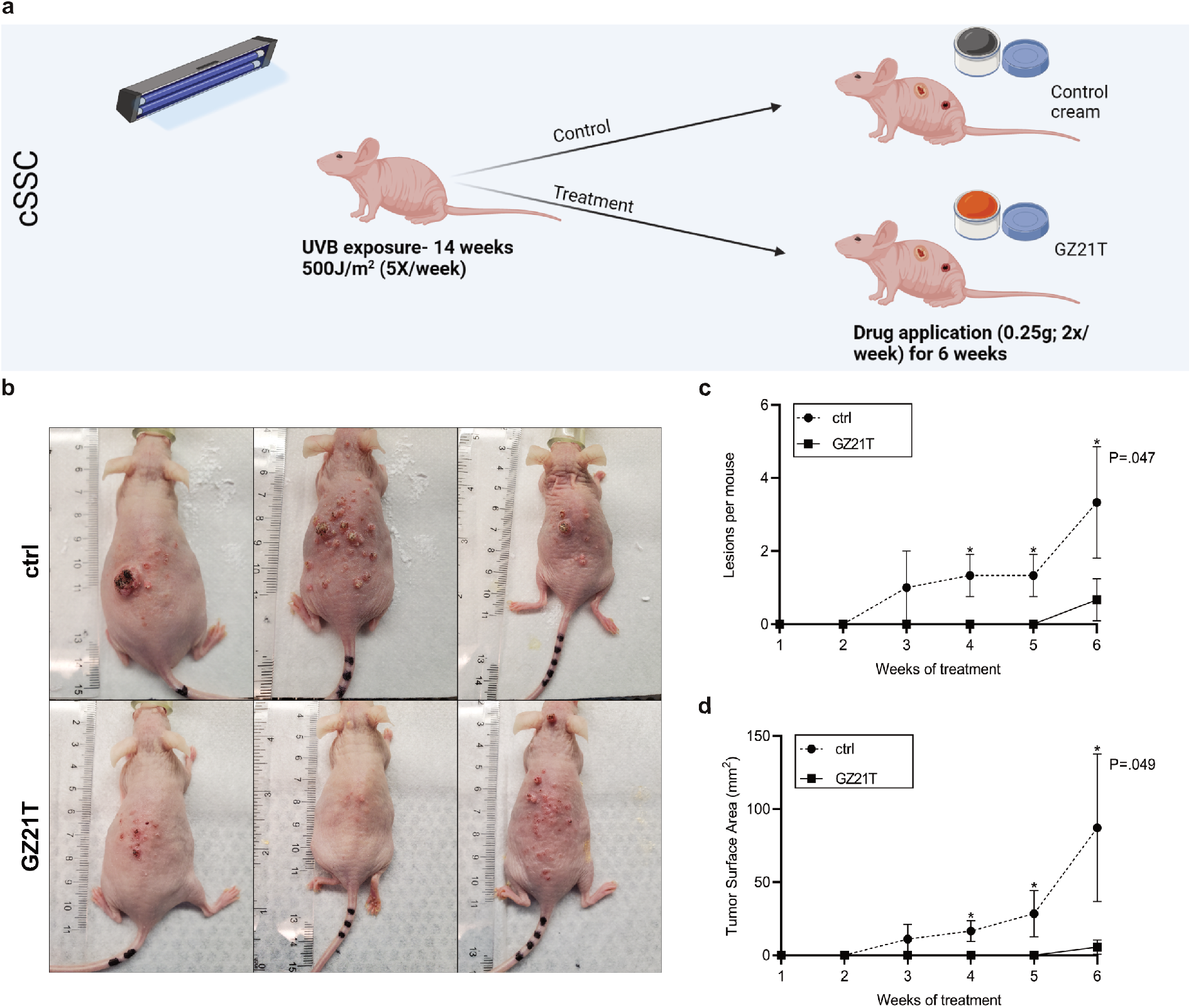
GZ21T inhibits the progression of actinic keratoses to cutaneous squamous cell carcinoma. A) To induce cSCC lesions, mice were irradiated with 500 J/m^2^ five days per week for 14 weeks and were treated with 0.25 g GZ21T or control cream two times per week for six weeks. B) Representative images of control and GZ21T treated mice after 6 weeks of treatment. C) Count and D) surface area of lesions greater than 3 mm in diameter. * P<0.05; cSCC, cutaneous squamous cell carcinoma; UVB, ultraviolet B; ctrl, control.

### GZ21T is cytoprotective to skin exposed to short-term UV radiation

We next evaluated the immediate histologic effects of GZ21T treatment with short-term, high-intensity UV exposure designed to mimic a sunny vacation (Figure 6A) (Mintie et al., 2020). Representative H&E images at the end of the UVB exposure period are shown in Figure 6B. Histological analysis revealed a significant decrease in mean epidermal thickness in the treated group compared to control mice (103.9±9.05 *μ*m vs. 157.0±15.32 *μ*m, *p*=.018; Figure 6C). We then assessed the expression of the proliferation marker Ki67 by immunohistochemistry (IHC) (Figure 6D). We found a significant decrease in the number of Ki67 positive cells in the epidermis of treated mice compared to the control group (79.6±12.8/HPF vs. 100.02±11.8/HPF, *p*=.042; Figure 6E). Finally, we conducted GSVA for pathways related to cell cycle progression in NL tissue from control and treated mice. This analysis showed suppression of G1/S transition (*p*=.04), mitotic DNA replication (*p*=.01), G2/M transition (*p*=.021), mitotic spindle assembly (*p*=.049), anaphase/metaphase transition (*p*=.02), and cytokinesis (*p*=.021) following GZ21T treatment (Figure 6F).

**Figure 6:**
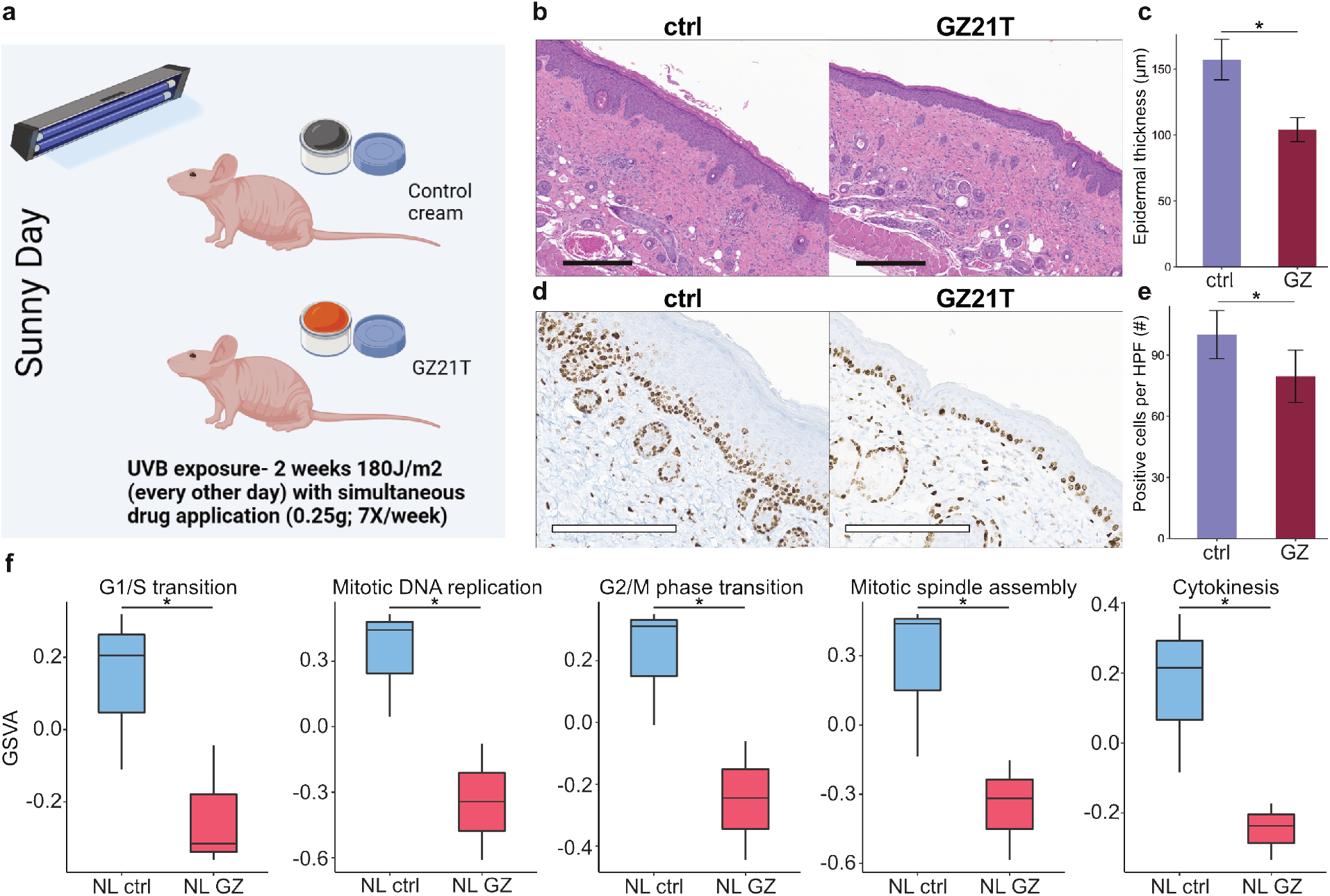
GZ21T exerts cytoprotective effects with short-term UV exposure. A) To assess the histological effects of GZ21T with short-term UV exposure, mice were irradiated with 180 J/m^2^ each day for two weeks with simultaneous application of 0.25 g GZ21T or control cream. B) Representative H&E images of control and GZ21T treated mice after two weeks of UV exposure and treatment. Black boxes represent 300 μm. C) Average epidermal thickness in control and GZ21T treated mice. D) Representative images of Ki67 staining in control and GZ21T treated mice after two weeks of UV exposure and treatment. White boxes represent 200 μm. E) Quantification of Ki67 staining in control and GZ21T treated mice. F) GSVA scores for pathways related to cell cycle progression in non-lesional samples of control and GZ21T treated mice. * P<0.05; UVB, ultraviolet B; ctrl, control; HPF, high powered field; GSVA, gene set variation analysis; NL, non-lesional; GZ, GZ21T.

## DISCUSSION

We demonstrate that topical GZ21T inhibits AK growth in a murine model of skin carcinogenesis. Mechanistically, GZ21T downregulates pathways involved in DNA synthesis, protein synthesis, metabolism, and cell cycle progression. Our results also show that GZ21T prevents the progression of AK to cSCC and exerts cytoprotective activity with short-term UV exposure.

While previous studies have utilized oral administration or intra-tumoral injection of GZ17-6.02, this is the first study to demonstrate its efficacy in a topical formulation. It is thought that topical administration of curcumin may have superior efficacy compared to oral dosing as the compound is susceptible to a large first pass metabolism resulting in less than 1% of oral curcumin entering the plasma (Helson, 2013). The topical formulation, GZ21T, can circumvent this drawback, as topical medications do not require significant systemic concentrations to reach the skin. Taken together, our findings suggest that the combination of a topical route of administration and the inherent synergism among components of GZ21T makes it an effective drug in the treatment of cutaneous disorders.

In the murine model of skin carcinogenesis, GZ21T was found to be a potent inhibitor of AK growth through its effect on multiple pathways and targets. Transcriptomic analysis of skin from untreated animals showed that progression from PL tissue to AK was accompanied by increases in ribosome, spliceosome, DNA replication, cell cycle, proteasome, and nucleotide excision repair related processes. These pathways were downregulated by GZ21T treatment.

GZ21T also suppressed several pathways previously implicated in the pathogenesis of AK and SCC, including MAPK, Pi3K-Akt, HIF-1α, Wnt, insulin, and ErbB signaling (Thomson et al., 2021). The MAPK pathway plays a crucial role in promoting the growth and survival of various malignancies (Fang & Richardson, 2005; Santarpia et al., 2012), and several studies suggest that the MAPK pathway may play a pivotal role in the transition from AK to SCC (Lambert et al., 2014; Zhang et al., 2018). Similarly, both Pi3K-Akt and insulin signaling pathways are frequently mutated in human cancers and serve to promote cell growth, survival, and metabolic processes by disconnecting the cell from control by exogenous growth signals (Hoxhaj & Manning, 2020; Malaguarnera & Belfiore, 2011). Aberrant insulin signaling may also promote a stem cell-like phenotype in malignant cells allowing for self-renewal and propagation of these tissues (Hoxhaj & Manning, 2020; Malaguarnera & Belfiore, 2011). Studies have shown that several key Pi3K-Akt and insulin-like-growth-factor receptor proteins are over-activated in AK and become progressively more dysregulated with progression to cSCC, highlighting the role of these pathways in UV-induced skin carcinogenesis (Einspahr et al., 2012; Thomson et al., 2021). HIF-1α is a transcription factor that is activated by intra-tumoral hypoxia to promote angiogenesis, glucose metabolism, migration, and invasion (Balamurugan, 2016). Additionally, overexpression of this protein has been associated with increased metastasis and higher mortality rates in several human cancers (Zhong et al., 1999; Zhou et al., 2006). The role of HIF-1α in AK specifically has been exemplified by murine knock-out models for the protein. Loss of HIF-1α results in decreased tumorigenesis as knock-out mice exhibit an increased rate of DNA repair processes, slowing the accumulation of carcinogenic mutations (Mahfouf et al., 2019). Aberrant Wnt signaling has also been shown to be a major factor in cSCC development and progression as it allows the transcription factor β-catenin to escape proteasomal degradation to promote a stem-like phenotype and hyperproliferation of keratinocytes (Lang et al., 2019). Finally, ErbB (EGFR) mutations are common in epithelial cancers, as they provide a growth stimulus to promote self-renewal and propagation of malignant tissues (Ciardiello & Tortora, 2008). Prior studies have shown that numeric aberrations in EGFR expression may be present in up to 52% of AKs and 77% of cSCC (Toll et al., 2010). Interestingly, these pathways were not altered in non-lesional skin, suggesting that GZ21T may preferentially suppress their signaling in AK lesions while sparing healthy tissue.

At the protein level, NL tissue was characterized by upregulation of the T-cell inhibitory protein PD-L1 (Ghosh et al., 2021) and glutaminase, a metabolic enzyme that promotes tumorigenesis (Wang et al., 2020). Progression from PL to AK resulted in increased expression of pAkt1 S473, eEF2K, and LCN2 with suppression of RPA32 and FANCD2. These changes were reversed with GZ21T treatment. The findings here are in line with previous studies showing that GZ17-6.02 treatment enhances the efficacy of anti-PD1 checkpoint inhibitors (Booth et al., 2020). pAkt1 S473 is increased in several head and neck squamous cell carcinomas (Islam et al., 2014) and promotes aberrant proliferation and survival of AK through the Pi3K-Akt pathway. Both eEF2K and LCN2 have been found to promote migration and invasion of cancer cells in esophageal squamous cell carcinomas (Du et al., 2015; Islam et al., 2014; Zhu et al., 2015) and may support the survival of neoplastic tissue under conditions of nutrient deprivation (Temme & Asquith, 2021). Meanwhile, RPA32 and FANCD2 play a key role in DNA Repair and DNA damage checkpoint responses (Alcón et al., 2020; Zou et al., 2006). Their upregulation suggests that GZ21T suppresses AK development by reducing the accumulation of carcinogenic mutations following UVB exposure. Finally, we found that GZ21T upregulates AMPK, a protein that promotes autophagy by phosphorylating proteins within the ULK1 and mTORC1 complexes (Li & Chen, 2019). This finding corroborates prior studies demonstrating that GZ17-6.02 promotes the death of AK cells by upregulating the autophagic flux (West et al., 2022). Taken together, the findings from transcriptomic and proteomic analyses indicate that GZ21T affects a wide range of cellular targets, and its ability to inhibit AK growth is likely multifactorial.

Finally, our *in vivo* data also demonstrate a robust inhibition in the progression of lesions greater than 3 mm in diameter and a reduction in epidermal thickness and number of Ki67 positive cells with short-term UV exposure. These findings are further supported by GSVA demonstrating suppression of several cell cycle pathways following GZ21T treatment. Increased keratinocyte proliferation is an adaptive response to UV damage that may eventually aid in the accumulation of mutations and progression to AK (Marionnet et al., 2014). As such, the findings here suggest that GZ21T exerts a cytoprotective effect in response to short term UV-exposure leading to a decrease in keratinocyte proliferation in response to UV damage.

Limitations of this study include its small sample size and testing only in mice. Future studies should be conducted to test GZ21T’s efficacy in other models and humans. In conclusion, GZ21T inhibits the growth of AK by modulating key drivers of multiple pathways such as MAPK, Pi3K-Akt, Wnt, and ErbB signaling and represents a promising therapeutic option for the treatment of AK.

## MATERIALS AND METHODS

### Ethical statement

All animal protocols used in this study were approved and performed in accordance with guidelines set forth by the Duke University Institutional Animal Care & Use Committee (IACUC) under protocol number A155-20-07.

### Animal experiments

For induction of AK lesions, female SKH1 mice (Charles River Laboratories, Wilmington, MA) between 6-8 weeks of age were irradiated with 500 J/m^2^ (the minimum erythemal dose (Pillon et al., 2017)) ultraviolet B five days per week for ten weeks, using a Phillips TL 40W12 RS SLV/25 fluorescent tube (Phillips, #928011301230) mounted 25 cm above the floor of a dedicated enclosure. Power output was measured using a PM100D power meter and S120VC power sensor (Thor Labs, #S120VC) and was used to calculate exposure time. Exposure times were internally re-calibrated on a weekly basis to account for temporal power degradation. Mice were monitored daily for signs of cutaneous injury. After the ten-week irradiation period, mice were randomized into control and treatment groups. Mice in the control group were treated with 0.25 g topical placebo cream five days per week, while mice in the treatment group received 0.25 g topical GZ21T five days per week. Mice were treated for eight weeks. Lesions were measured using digital calipers, and tumor burden was quantified as total lesion count and surface area occupied by tumor.

To evaluate the ability of GZ21T to prevent the progression of AK to cSCC, female SKH1 mice between 6-8 weeks of age were irradiated with 500 J/m^2^ UVB five days per week for 14 weeks. Once lesions appeared, mice were randomized into control and treatment groups and received 0.25 g topical control cream or GZ21T twice weekly for six weeks. The dosing frequency was reduced compared to the AK experiment to determine if GZ21T could exert anti-tumor activity with less frequent application. Tumor burden of lesions likely to be cSCC was assessed using the total count and surface area occupied by lesions greater than 3 mm in diameter, as lesions of this size are more likely to represent cSCC than AK (de Gruijl & van der Leun, 1991).

To assess the short-term histopathologic effects of GZ21T on UVB-induced epidermal proliferation, female SKH1 mice between 6-8 weeks of age were exposed to UVB (180 mJ/cm^2^) every other day for two weeks, which mimics sun exposure during a sunny vacation (Mintie et al 2020). Mice in the control group were treated with 0.25 g topical placebo cream daily, while mice in the treatment group were treated with 0.25 g topical GZ21T.

### Statistical analysis

Statistical analyses were conducted using GraphPad Prism 9.0 software. Comparisons between groups were made using Student’s *t*-tests. Differences with a p-value of < 0.05 were considered statistically significant. Details of mRNA sequencing, RPPA, histology, and IHC are provided in the Supplementary Materials and Methods.

## SUPPLEMENTAL MATERIALS AND METHODS

### mRNA sequencing

Lesional samples were collected from AK lesions, PL samples were taken from skin immediately adjacent to a lesional area, NL samples were collected from normal-appearing skin at least 5 cm away from a lesional area, and non-UV exposed samples were collected from the skin on the ventral surface. All skin samples were flash-frozen in liquid nitrogen and stored at −80°C. Total RNA was isolated using an RNAeasy plus kit (Qiagen, #74034) according to the manufacturer’s instructions. RNA-seq data were processed using the TrimGalore toolkit (https://www.bioinformatics.babraham.ac.uk/projects/trim_galore). Only reads that were 20nt or longer after trimming were kept for further analysis. Reads were mapped to the GRCh38v93 version of the human genome and transcriptome (Kersey et al., 2012) using the STAR RNA-seq alignment tool (Dobin et al., 2013). Reads were kept for subsequent analysis if they mapped to a single genomic location. Gene counts were compiled using the HTSeq tool (http://www.huber.embl.de/users/anders/HTSeq). Only genes with at least 10 reads in any given library were used in subsequent analyses. Normalization and differential expression were carried out using the DESeq2 (Love et al., 2014) Bioconductor (Huber et al., 2015) package with the R statistical programming environment (www.r-project.org). The false discovery rate was calculated to control for multiple hypothesis testing. GSEA was performed using the GSEA software (Mootha et al., 2003; Subramanian et al., 2005) with the Kyoto Encyclopedia of Genes and Genomes (KEGG) database as a reference. Gene set variation analysis was conducted with the GSVA R Bioconductor package using the R statistical programming environment (Hänzelmann et al., 2013). Differentially expressed genes were defined as coding genes with a log fold change > 1 or < −1 and a false discovery rate-adjusted *P*-value < 0.05.

### Reverse Phase Protein Array (RPPA)

Lesional, PL, NL, and non-UV exposed skin were collected from mice, flash-frozen in liquid nitrogen, and stored at −80°C prior to being sent to MD Anderson’s RPPA core facility for processing. KEGG functional enrichment analyses were performed on gene lists of proteins whose expression decreased with GZ21T treatment using enrichR (Chen et al., 2013; Kuleshov et al., 2016; Xie et al., 2021).

### Histological assessment

At the end of the *in vivo* experiments, mice were euthanized, and dorsal skin samples were fixed in 10% neutral-buffered formalin. Formalin-fixed tissues were paraffin-embedded, sectioned, and mounted on serial slides at the Research Immunohistochemistry Lab at Duke University School of Medicine. Slides were stained with hematoxylin and eosin for histological determination.

### Immunohistochemistry

Ki67 staining was performed using a rabbit Ki67 antibody (Abcam, #ab16667) at a 1:200 dilution with Discovery Antibody Diluent (Roche, #760-108). IHC tests were performed using the Ultra Discovery automated staining platform (Roche, #05987750001). Tissue sections were pretreated with Roche cell conditioning solution CC1 (Roche, #950-124) for 56 minutes and were subsequently incubated with rabbit monoclonal Ki67 for 60 minutes at 36°C. Rabbit IgG, substituted for the primary antibody, was used as a negative control. After binding of the primary antibody, Roche anti-rabbit HQ (Roche, #760-4815) was applied and incubated for 12 minutes, followed by a 12-minute incubation with anti-HQ HRP (Roche, #760-4820) for antigen detection. The IHC reaction was visualized with DAB chromogen and counterstained hematoxylin.

## DATA AVAILABILITY STATEMENT

The data that support the findings of this study are available from the corresponding author M.M.K., upon reasonable request.

## ORCIDs

Junwen Deng: http://orcid.org/0000-0001-8393-8915

Varsha Parthasarathy: http://orcid.org/0000-0002-1422-772X

## AUTHOR CONTRIBUTIONS

Conceptualization: ZAB, JC, CEW, SGK, MMK

Data Curation: ZAB, JC, GB, CD

Formal Analysis: ZAB, JC

Funding Acquisition: MMK

Investigation: ZAB, JC, GB, CD, CEW, SGK, MMK

Methodology: ZAB, JC, CEW, SGK, MMK

Project Administration: CEW, SGK, MMK

Resources: CEW, SGK, MMK

Software: ZAB, JC

Supervision: CEW, SGK, MMK

Validation: CEW, SGK, MMK

Visualization: ZAB, JC

Writing – Original Draft Preparation: ZAB, JC, MM, KL, CS, CEW, SGK, MMK

Writing – Review and Editing: ZAB, JC, GB, CD, MM, KL, CS, JA, RW, AP, AK, HC, SVR, WW, OOO, MPA, CEW, SGK, MMK

---

**Figure S1:**
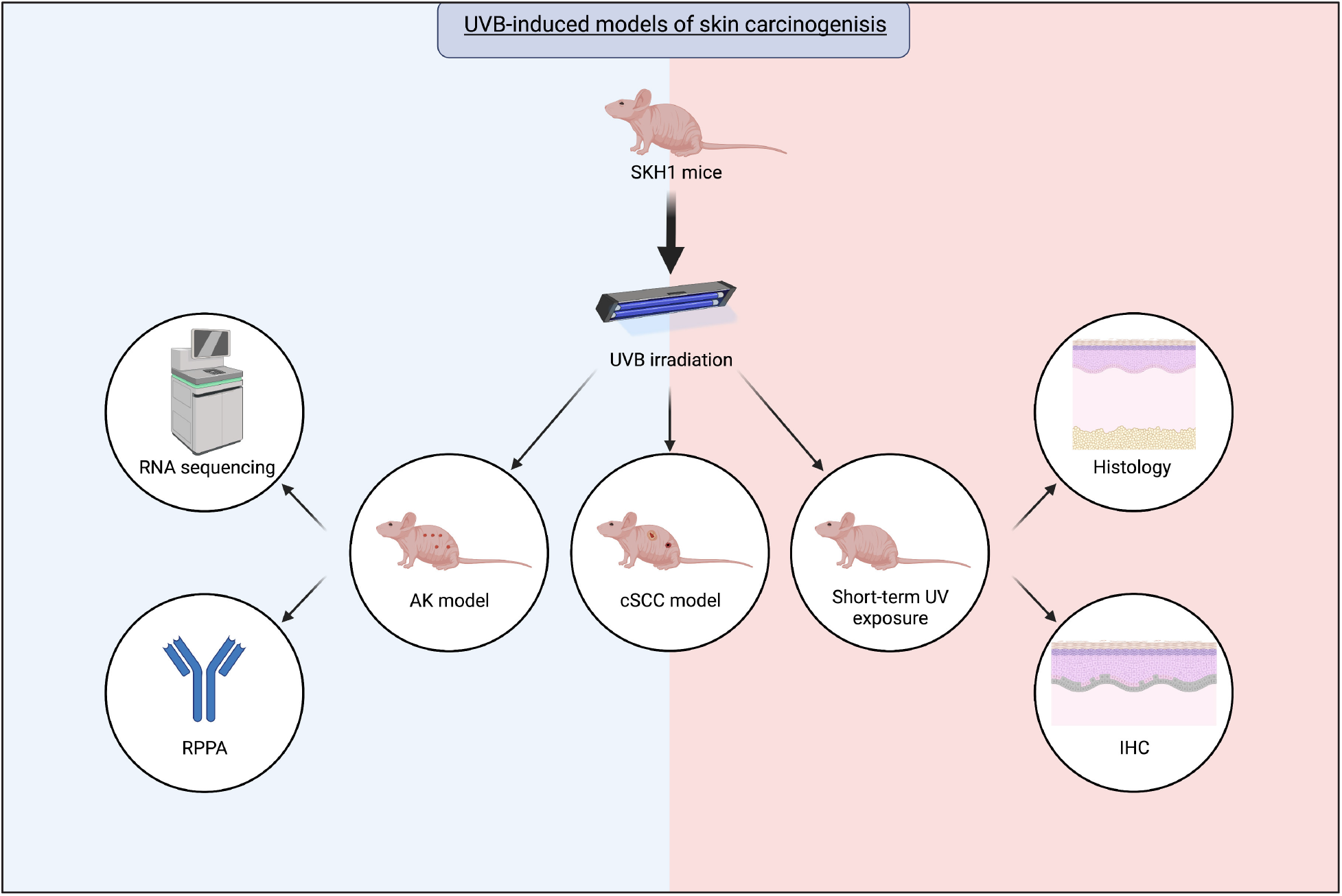
Overall study design. SKH1 mice were used for *in vivo* models of skin carcinogenesis. Varying UVB exposure protocols were used to evaluate the effects of GZ21T with a model of AK, a model of cSCC, and short-term UV exposure designed to mimic a sunny vacation. RNA sequencing and reverse phase protein arrays were performed on skin samples from the AK experiments, while histology and IHC were performed on skin from the short-term UV exposure experiments. UVB, ultraviolet B; RPPA, reverse phase protein arrays; AK, actinic keratosis; cSCC, cutaneous squamous cell carcinoma; IHC, immunohistochemistry.

